# Probabilistic Edge Inference of Gene Networks with Bayesian Markov Random Field Modelling

**DOI:** 10.1101/2022.07.30.501645

**Authors:** Yu-Jyun Huang, Rajarshi Mukherjee, Chuhsing Kate Hsiao

## Abstract

Gaussian graphical models (GGMs), also known as Gaussian Markov random field (MRF) models, are commonly used for gene regulatory network construction. Most current approaches to estimating network structure via GGMs can be categorized into a binary decision that determines if an edge exists through penalized optimization and a probabilistic approach that incorporates graph uncertainty. Analyses in the first category usually adopt the perspective of variable (edge) selection without consideration of probabilistic interpretation. Methods in the second group, particularly the Bayesian approach, often quantify the uncertainty in the network structure with a stochastic measure on the precision matrix. Nevertheless, these methods overlook the existence probability of an edge and its strength related to the dependence between nodes. This study simultaneously investigates the existence and intensity of edges for network structure learning. We propose a method that combines the Bayesian MRF model and conditional autoregressive model for the relationship between gene nodes. This analysis can evaluate the relative strength of the edges and further prioritize the edges of interest. Simulations and a glioblastoma cancer study were carried out to assess the proposed model’s performance and compare it with existing methods. The proposed approach shows stable performance and may identify novel structures with biological insights.

## 1. INTRODUCTION

Structural information analysis of multivariate data has attracted much attention in recent decades, especially in the biomedical research community (Chang et al., 2020; Wang et al., 2021). One common approach to describe the network structure of a group of genetic variables and the conditional dependence between them is the Markov random field (MRF), an undirected graphical model (Lauritzen, 1996; Drton and Maathuis, 2017). Examples include gene regulatory networks, brain connectivity networks, and microbial networks (Cai et al., 2019; Zhang et al., 2019; Huang et al., 2020). The Gaussian MRF, also known as the Gaussian graphical model (GGM), is a widely used multivariate distribution for gene regulatory networks. It assumes that the *p*-dimensional vector 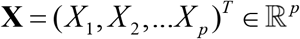 follows a multivariate normal distribution 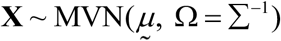 with *X_i_* denoting the gene expression value of the *i*-th gene node. A zero-entry in the precision matrix Ω corresponds to conditional independence and no connecting line between nodes. In other words, if the off-diagonal (*i, j*)-th element *ω_ij_* in Ω is zero, then the partial correlation 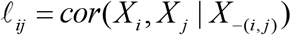 is zero; namely, the *X_i_* and *X_j_* are conditionally independent given the remaining variables, and there exists no edge between these paired nodes in the network. In other words, under GGM, the problem of network construction becomes the inference of a sparse precision matrix or the selection of non-zero partial correlation.

Recent work on inferring network structure with GGM can be categorized into two groups. Methods in the first group focus on determining if an edge exists between nodes using the idea of “covariance selection” coined by Dempster (1972). When *p* is large, these methods follow the principle of variable selection with a regularization procedure to complete the binary decision about whether *ω_ij_* or ℓ*_ij_* is zero. Various methods of this regularization approach have been developed that adopt different objective functions and/or *L*_1_ penalty, including neighborhood selection with lasso by Meinshausen and Buhlmann (2006), graphical lasso (Glasso) in Friedman et al. (2008), the space partial correlation estimation (SPACE) in Peng et al. (2009), and the constrained *l*_1_ minimization for inverse matrix estimation (CLIME) in Cai et al. (2011). These penalized optimization methods can be applied straightforwardly, but they are not designed to infer the intensity of edges or to interpret the dependence between nodes. Such information can be influential in biological experiments (Ni et al., 2020). If the inference, such as the estimation of the non-zero partial correlation, is based on a given network, then the network structure needs to be fixed first with one of the methods mentioned above. Therefore, this estimation procedure relies heavily on the choice of the selected network structure, which may cause concern about subsequent inference if the validity of this structure is in question.

Methods in the second group, usually under the Bayesian framework, explicitly adopt the uncertainty in the network graph. In this probabilistic graphical modeling, Wang and Pillai (2013) used a shrinkage prior, while both Wang and Li (2012) and Mohammadi and Wit (2015) incorporated the G-Wishart prior on the precision matrix. They proposed various tools, such as the double Metropolis-Hasting algorithm and birth-death Markov chain Monte Carlo methods, to enhance computational efficiency. Their analysis can provide a posterior probability for each candidate graph and a posterior inclusion probability for each edge. The inclusion probability, in this case, can be a good indication of its existence, but the strength of the edge is not considered in the computation. One solution would be to average the estimates of precision matrices in an element-wise way and weight by the posterior probability of the matrix and the corresponding candidate graph. For instance, the BDgraph in Mohammadi and Wit (2015) can be utilized to perform this analysis. However, the computational burden in these procedures is already fairly heavy due to a large number of nodes and the even more significant number of candidate graphs. If the strength of the edge is to be inferred further in these analyses, challenges arise from the increased computational load.

The rationale of this research is twofold. First, an informative metric to quantify the strength of an edge is needed, which can provide more information beyond its existence. This is crucial when decoding the interplay between nodes or prioritizing intervention in a gene regulatory network. Second, since most genes do not work alone, the strength or intensity of the relationship between any two nodes should account for the presence of other genes when learning the network structure of a given set of genetic nodes. In this study, we start with the Bayesian MRF combining the conditional autoregressive (CAR) model to estimate the strength of the edge and its existence probability. Under the Gaussian CAR model, the conditional mean *E*(*X_j_* | *X*_(−*j*)_) is expressed as 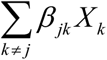 for *j* = 1, 2, …, *p*, where 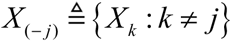 represents the set containing all variables except *X_j_*. Following Besag (1974) and Besag and Kooperberg (1995), the coefficient *β_jk_* is a function of elements in the precision matrix Ω, and is connected to the partial correlation ℓ*_jk_* between *X_j_* and *X_k_*. That is, the *β_jk_* can be used to characterize the strength of dependence between these two genes. In addition, the Spike-and-Slab Lasso (SSL) prior proposed by Ročková and George (2018) is adopted for *β_jk_*. Then, the regularization procedure on these *β_jk_* ’s functions similarly to the “covariance selection” procedure in previous literature and provides a direct and intuitive interpretation of the intensity and relationship between nodes.

The rest of this article is organized as follows. The rationale and complete model of the Bayesian MRF and the implementation of prior knowledge are introduced in Section 2. In Section 3, extensive simulation studies are conducted to demonstrate the performance of the proposed model and comparison with other state-of-the-art methods. In Section 4, the proposed model is applied to a glioblastoma study with gene expression values from TCGA (Hutter and Zenklusen, 2018). Some biologically relevant findings will be highlighted. We then conclude with a discussion.

## 2. BAYESIAN MARKOV RANDOM FIELD MODEL

### 2.1 Learning network structure

To introduce the proposed Bayesian Markov Random field (BMRF) model, we first let the *n* × *p* matrix **X** represent the observed gene expression values of the *p* genes from the *n* subjects, where *x_ij_* is the expression value of the *j*-th gene (*j* = 1,2, …, *p*) from the *i*-th subject (*i* = 1,2, …, *n*). Without loss of generality, the values across subjects per gene are standardized so that *E*(*X_j_*) = 0 and *Var*(*X_j_*) = 1. Under GGM, the *p*-dimensional random vector (*X*_1_, *X*_2_, …, *X_p_*)*^T^* follows a multivariate normal distribution (MVN) with the following conditional distribution (Besag 1974),

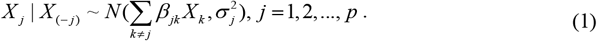

Following Besag (1974) and Besag and Kooperberg (1995), the coefficients can be expressed as 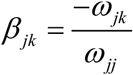 if *j* ≠ *k*. This is related to the partial correlation ℓ*_jk_* between *X_j_* and *X_k_* where 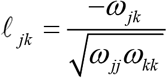. When the diagonal elements in Ω are equal, then *β_jk_* = *β_kj_* and the underlying coefficients in the CAR model can be expressed as 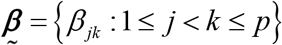 where 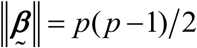 is the number of unknown parameters to be estimated. Moreover, when *β_jk_* = 0, the corresponding ℓ_*jk*_ = 0, implying no edge between two gene nodes. These properties provide two advantages in supporting *β_jk_* as promising candidates in inferring the network structure. First, the selection of non-zero elements of 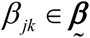 is equivalent to the decision of the existence of the edge. Second, the magnitude of these coefficients can quantify the relative intensity of partial correlation between nodes, especially when a direct estimate of the correlation is not straightforward due to the curse of dimensionality.

This CAR model is more general than those used in spatial statistics, where only neighboring “areas” are included in the mean structure. Here all genetic nodes are included first as a fully connected model. Then the procedures and computations below will decide which *β_jk_* remain and how strong the evidence is. In addition, this conditional distribution is also similar to node-wise regression where constraints are imposed to ensure symmetry in the *β_jk_*’s (Ha et al. 2021).

### 2.2 Spike-and-Slab Lasso prior: probabilistic estimation of edge

For the inference of *β_jk_*, we consider the Spike-and-Slab Lasso (SSL) prior (Rockova and George, 2018),

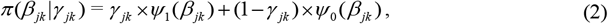

where the slab distribution 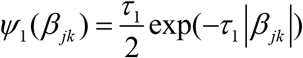 and the spike *ψ*_0_ (*β_jk_*) are both double exponential with a small *τ*_1_ and large *τ*_0_, respectively.

In contrast to the previously mentioned penalized optimization methods for variable selection, where the estimated effect size is biased, the SSL prior is considered a fundamental variable selection tool in the Bayesian framework for sparse models (George and McCulloch, 1993). The binary *γ_jk_* takes the value of 1 if *β_jk_* generates a large effect. Therefore, the marginal posterior probability of *γ_jk_* = 1 represents the posterior probability of the edge existence and would be a good indication of an existent edge. In addition, the SSL prior is flexible because it allows the shrinkage effects to vary among different edges. For instance, a substantial shrinkage penalty can be deployed for those edges with weak partial correlation, while for those with strong partial correlation, a non-shrinkage effect can be considered. By adopting the SSL prior, we can select the influential edges and perform statistical inference with *β_jk_*. The BMRF model specification is completed with a Bernoulli prior for *γ_jk_*, *γ_jk_* ~ *Ber*(*p_jk_*), where *p_jk_* follows a conjugate beta distribution.

### 2.3 Computation

Since the posterior distributions of *γ_jk_* and *β_jk_* are the bases of the probabilistic inference, one can obtain the posterior samples of *γ_jk_* and *β_jk_* with Markov chain Monte Carlo (MCMC) methods implemented in any standard Bayesian software. In the following simulation studies and applications, the R package *R2OpenBUGS* is used to carry out the computations.

When the number of gene nodes is large, the number of possible edges and parameters increases rapidly. Fortunately, most genetic networks/pathways are sparse. For instance, the sparsity of the signaling pathway networks in KEGG ranges between 5% and 10%. Liu et al. (2009), Zhao et al. (2012), and Mohammadi and Wit (2015) have adopted similar values in their simulation studies. Such *a priori* information can be utilized in a *p* × *p* adjacency matrix *G\**, where elements *g_jk_* = 1 if two genes *X_j_* and *X_k_* are known biologically to be associated and *g_jk_* = 0 otherwise. By imposing the matrix of domain knowledge *G\** on 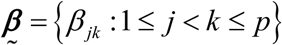, one can save computational cost from estimating the edges known to be non-existent. Similarly, another *p* × *p* adjacency matrix *M\** can be introduced to contain elements *m_jk_* = 1 if the corresponding interrelation is of interest to particular experts. This would force the inclusion of the edge in the network, yet the flexibility remains when later inference does not favor its existence. Inclusion of these two matrices and the distribution of *p_jk_* can account for all the cases described here. For example, this matrix *M\** can be derived first and the data-driven prior on *γ_jk_* can be further established. The BMRF with this setup will be denoted as BMRF.P in later sections.

## 3. SIMULATION STUDIES

For performance evaluation and comparison with existing methods, three types of network graph are considered in the simulation studies: the random network (M1), random scale-free network (M2), and fixed network structure (M3). In M1, edges are considered exchangeable, and all nodes in a network are equally important. The scale-free network in M2 is commonly adopted for genetic pathways, where the edges are not exchangeable because hub nodes may exist in the network. These two are designed to compare with the traditional approach of variable selection, where only the number of true edges successfully detected is of concern. While in M3, with a fixed and known structure, further comparison between the inclusion probability in previous Bayesian methods and the existence probability in current BMRF can be carried out, and the strength of edge is demonstrated. In other words, in M3, in addition to the number of true edges successfully detected, both the probability of existence and strength of edges will be emphasized.

### 3.1 Settings

In the *random network* setting M1, the GGM is generated with the following steps, similar to the procedures in Fan et al. (2009), Peng et al. (2009), and Yin and Li (2011).

1. Set up the network sparsity *S*, 0 ≤ *S* ≤ 1.
2. Construct the true network *E* by randomly sampling the Bernoulli *e_ij_* with probability *S*. If *e_ij_* = 1, then there is an edge between the node *i* and *j*, and 0 otherwise.
3. Generate the precision matrix 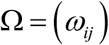 according to *E* by

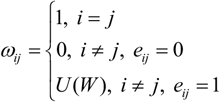

where 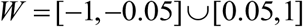 and *U*(.) denotes the uniform distribution.
4. To assure the positive definiteness of Ω, each off-diagonal *ω_ij_* in Ω is replaced by the original *ω_ij_* divided by 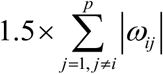.
5. Average the rescaled matrix calculated in (4) with its transpose matrix to ensure symmetry. The values of the nodes are generated from a multivariate normal distribution (MVN) with a zero mean vector and the precision matrix.

Note that different combinations of *p* and *S* have been considered, denoted as M1.1 for *p* = 25, *S* = 0.05, M1.2 for *p* = 25, *S* = 0.10, M1.3 for *p* = 50, *S* = 0.05, and M1.4 for *p* = 50, *S* = 0.10. The number of edges in each network is about 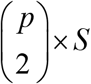.

In the *random scale-free network* setting M2, the R package *huge* was used to generate the scale-free networks. Two settings (M2.1) *p* = 25 and (M2.2) *p* = 50 were considered. The average number of edges in the scale-free network is *p* – 1.

In M3, the *fixed network structure* setting, a scale-free network graph containing 50 nodes and 49 edges was selected, and the node values were generated with the *huge* package with the partial correlation in the network set at −0.216.

For all stimulations, the hyper-parameters were specified as *τ*_1_ = 2 and *τ*_0_ = 20, the sample size was *n* = 250, and the number of replications in each setting was 100. More detailed information, including the network sparsity and number of true edges, is summarized in the Supporting Information Table S1. For the proposed BMRF, the corresponding edge is selected for the network if the posterior probability of *γ_jk_* = 1 is greater than 0.5. This choice is used in simulation studies when comparing different regularized methods for variable selection.

### 3.2 Comparison and criteria

The proposed BMRF model was compared with M&B (Meinshausen and Buhlmann, 2006), Glasso (Firedman et al., 2008), SPACE (Peng et al., 2009), and CLIME (Cai et al., 2011), as well as with the Bayesian approach BDgraph (Mohammadi and Wit, 2015) using the Bayesian model averaging procedure (denoted as BD_BMA) or the Maximum a posterior probability procedure (BD_MAP). M&B and Glasso were performed with the R package *huge*, and the tuning parameter used in these two methods was chosen through the rotation information criterion (ric). The SPACE approach was performed with the R package *space* with the tuning parameter set by default. The package *flare* was used for the estimator CLIME with tuning parameters obtained by 5-fold cross-validation. The R package *BDgraph* was used for BDgraph.

Several criteria were used to compare performance, including the total number of true positives (TP), the sensitivity (SEN), the specificity (SPE), the false discovery rate (FDR), the Matthew correlation coefficient (MCC), and the F1-score (F1). These quantities are calculated based on TP and the total number of false negatives (FN), where TP is defined as the total number of true edges that were successfully identified, and FN as the total number of true edges that failed to be detected.

### 3.3 Implementations

When handling a large set of gene nodes with BMRF, we recommend two modeling strategies, one with a non-informative prior and the other with a data-driven prior. The former is denoted as BMRF.O, corresponding to the prior distribution *γ_jk_* ~ *Ber*(*p_jk_*) with *p_jk_* from a beta distribution with mean 0.5. The latter, denoted as BMRF.P, models the network edges with *p_jk_* ~ *Beta*(*α\**, *β\**), an informative prior with a mean larger than 0.5 if 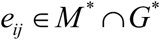, or *p_jk_* ~ *Beta*(*α^†^*, *β*^†^), a non-informative prior with a mean around 0.5. As stated earlier, the matrices *M\** and *G\** can be elicited by experts, with domain knowledge, with a screening scheme based on sparsity or sample correlation, or with SPACE proposed in Peng et al. (2009), which outperforms other methods when dealing with a scale-free network structure. In the following analysis, the matrix *G*\* containing the edges corresponding to the largest 10% absolute sample correlations was determined first when the network sparsity was set at 0.05 (or the top 15% if set at 0.10). For the matrix *M*\*, we incorporated the information from SPACE to accelerate the computational efficiency. The mean of the informative prior *p_jk_* ~ *Beta*(*α\**, *β*\*) was set at 0.8.

### 3.5 Results

#### 3.5.1 Random network (M1 and M2)

To compare performance, Table 1 lists the evaluation criteria under settings M1.1, M1.3, and M2.2. Specifically, the F1-score is displayed in Figure 1. A quick look indicates that the proposed BMRF.O and BMRF.P (pink color) are among the best. Judging from the F1-score and from the MCC in Table 1, M&B, SPACE, BMRF.O, and BMRF.P performed better than the others. Among these four, M&B had both a slightly larger F1-score and MCC, and it achieved the smallest FDR. However, its sensitivity and TP were not as good as SPACE and BMRF.P, while BMRF.P outperformed SPACE on these two criteria. The performances under other settings are displayed in the Supporting Information Table S2. Among the Bayesian methods, BD_BMA and BD_MAP tended to identify more edges, leading to larger TP and SEN but lower F1 and MCC. Consequently, these two often produce a larger FDR. BD_MAP was usually the worst in this regard due to the lack of consideration of model uncertainty.

**Table 1.**
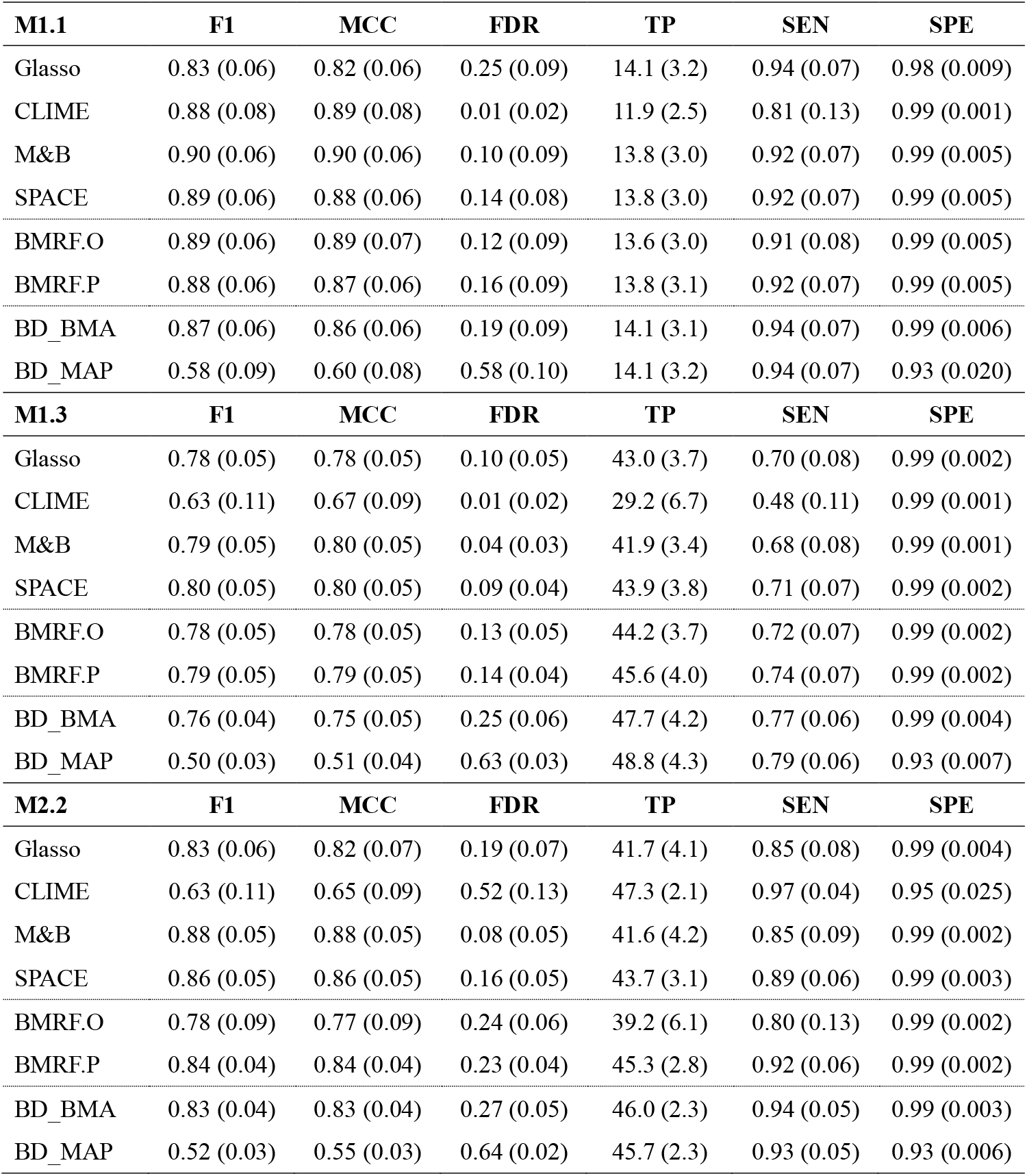
Values of evaluation criteria under simulation settings M1.1, M1.3, and M2.2. Each value is the average of 100 replications with standard error (SE) in parentheses.

| **M1.1** | **F1** | **MCC** | **FDR** | **TP** | **SEN** | **SPE** |
| --- | --- | --- | --- | --- | --- | --- |
| Glasso | 0.83 (0.06) | 0.82 (0.06) | 0.25 (0.09) | 14.1 (3.2) | 0.94 (0.07) | 0.98 (0.009) |
| CLIME | 0.88 (0.08) | 0.89 (0.08) | 0.01 (0.02) | 11.9 (2.5) | 0.81 (0.13) | 0.99 (0.001) |
| M&B | 0.90 (0.06) | 0.90 (0.06) | 0.10 (0.09) | 13.8 (3.0) | 0.92 (0.07) | 0.99 (0.005) |
| SPACE | 0.89 (0.06) | 0.88 (0.06) | 0.14 (0.08) | 13.8 (3.0) | 0.92 (0.07) | 0.99 (0.005) |
| BMRF.O | 0.89 (0.06) | 0.89 (0.07) | 0.12 (0.09) | 13.6 (3.0) | 0.91 (0.08) | 0.99 (0.005) |
| BMRF.P | 0.88 (0.06) | 0.87 (0.06) | 0.16 (0.09) | 13.8 (3.1) | 0.92 (0.07) | 0.99 (0.005) |
| BD_BMA | 0.87 (0.06) | 0.86 (0.06) | 0.19 (0.09) | 14.1 (3.1) | 0.94 (0.07) | 0.99 (0.006) |
| BD_MAP | 0.58 (0.09) | 0.60 (0.08) | 0.58 (0.10) | 14.1 (3.2) | 0.94 (0.07) | 0.93 (0.020) |
| **M1.3** | **F1** | **MCC** | **FDR** | **TP** | **SEN** | **SPE** |
| Glasso | 0.78 (0.05) | 0.78 (0.05) | 0.10 (0.05) | 43.0 (3.7) | 0.70 (0.08) | 0.99 (0.002) |
| CLIME | 0.63 (0.11) | 0.67 (0.09) | 0.01 (0.02) | 29.2 (6.7) | 0.48 (0.11) | 0.99 (0.001) |
| M&B | 0.79 (0.05) | 0.80 (0.05) | 0.04 (0.03) | 41.9 (3.4) | 0.68 (0.08) | 0.99 (0.001) |
| SPACE | 0.80 (0.05) | 0.80 (0.05) | 0.09 (0.04) | 43.9 (3.8) | 0.71 (0.07) | 0.99 (0.002) |
| BMRF.O | 0.78 (0.05) | 0.78 (0.05) | 0.13 (0.05) | 44.2 (3.7) | 0.72 (0.07) | 0.99 (0.002) |
| BMRF.P | 0.79 (0.05) | 0.79 (0.05) | 0.14 (0.04) | 45.6 (4.0) | 0.74 (0.07) | 0.99 (0.002) |
| BD_BMA | 0.76 (0.04) | 0.75 (0.05) | 0.25 (0.06) | 47.7 (4.2) | 0.77 (0.06) | 0.99 (0.004) |
| BD_MAP | 0.50 (0.03) | 0.51 (0.04) | 0.63 (0.03) | 48.8 (4.3) | 0.79 (0.06) | 0.93 (0.007) |
| **M2.2** | **F1** | **MCC** | **FDR** | **TP** | **SEN** | **SPE** |
| Glasso | 0.83 (0.06) | 0.82 (0.07) | 0.19 (0.07) | 41.7 (4.1) | 0.85 (0.08) | 0.99 (0.004) |
| CLIME | 0.63 (0.11) | 0.65 (0.09) | 0.52 (0.13) | 47.3 (2.1) | 0.97 (0.04) | 0.95 (0.025) |
| M&B | 0.88 (0.05) | 0.88 (0.05) | 0.08 (0.05) | 41.6 (4.2) | 0.85 (0.09) | 0.99 (0.002) |
| SPACE | 0.86 (0.05) | 0.86 (0.05) | 0.16 (0.05) | 43.7 (3.1) | 0.89 (0.06) | 0.99 (0.003) |
| BMRF.O | 0.78 (0.09) | 0.77 (0.09) | 0.24 (0.06) | 39.2 (6.1) | 0.80 (0.13) | 0.99 (0.002) |
| BMRF.P | 0.84 (0.04) | 0.84 (0.04) | 0.23 (0.04) | 45.3 (2.8) | 0.92 (0.06) | 0.99 (0.002) |
| BD_BMA | 0.83 (0.04) | 0.83 (0.04) | 0.27 (0.05) | 46.0 (2.3) | 0.94 (0.05) | 0.99 (0.003) |
| BD_MAP | 0.52 (0.03) | 0.55 (0.03) | 0.64 (0.02) | 45.7 (2.3) | 0.93 (0.05) | 0.93 (0.006) |

**Figure 1.**
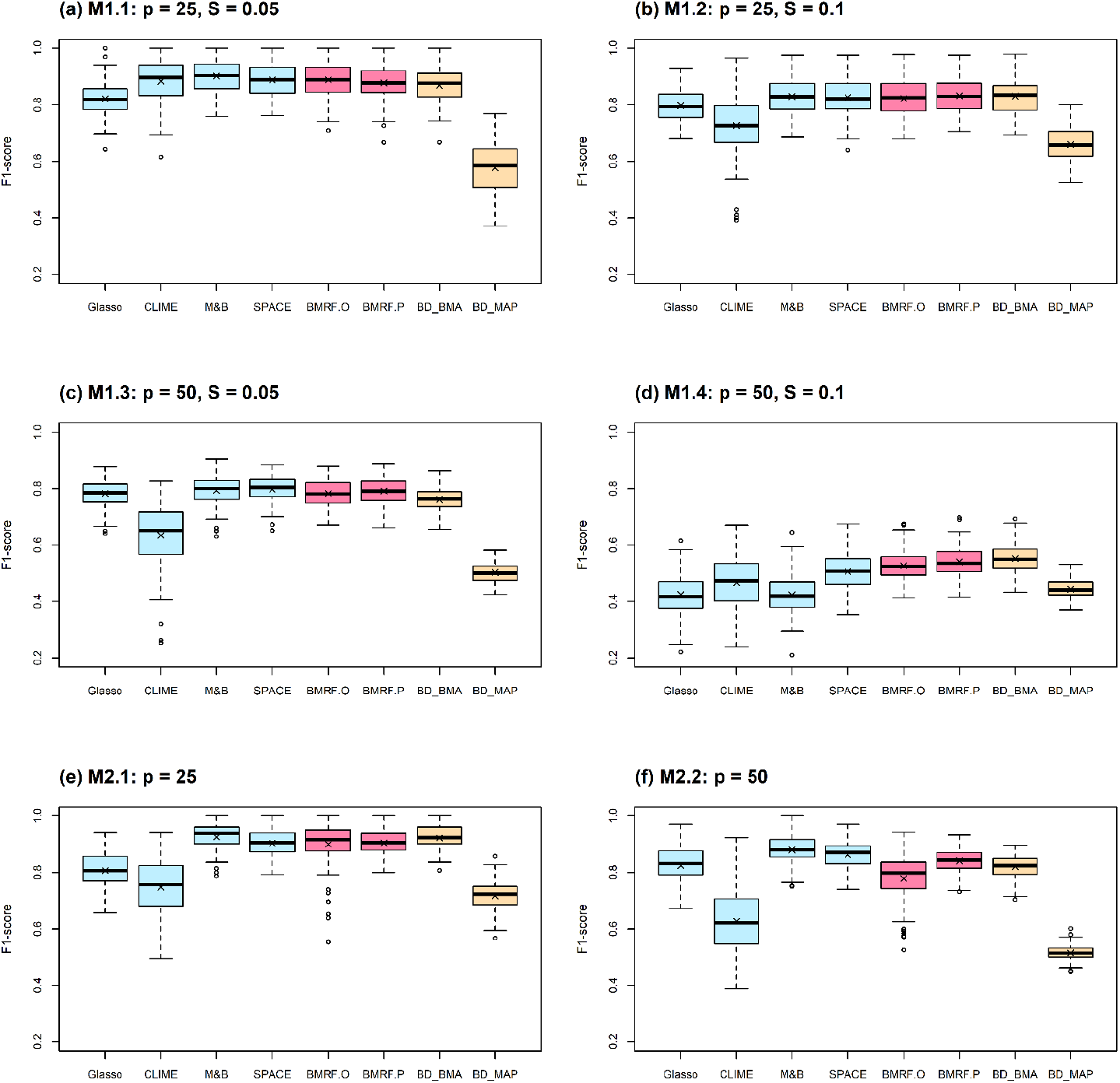
Boxplots of F1-scores from 100 replications under each method. Each subfigure contains a different combination of *p* and *S*. Blue boxplots correspond to the penalized methods, pink to the two BMRF models, and yellow to the two BDgraph tools.

In setting M1.4, when the number of nodes *p* is as large as 50 and the sparsity *S* = 0.1, most methods performed unsatisfactorily (Figure 1 (d)). However, the Bayesian approaches—both BMRF and BDgraph—seemed to be less sensitive to the increase in *p* and *S*. This is because these methods tend to identify more edges when compared with the frequentist approach to variable selection, therefore leading to a higher F1-score. These results highlight the advantages of probabilistic inference on the conditional dependence in network analysis, in contrast to the detection of whether or not the edge exists.

#### 3.5.2 Fixed network (M3)

In setting M3, a fixed network structure with two hub nodes was determined first, as shown in Figure 2 (a), and then the node values were generated from MVN. The numbers of edges connecting to the two hubs, Node-2 and Node-4, are 14 and 17, respectively. Various methods were then applied to infer the network structure. Across 100 replications, the average number of edges estimated by each method is listed in Table 2. Three methods, M&B, SPACE, and BMRF.P, performed the best, with BMRF.P being slightly better with a smaller standard error. The same pattern is observed in the F1-score shown in Figure 2 (b). The values of the other criteria are summarized in the Supporting Information Tables S3. Among these competing methods, only three, BMRF.O, BMRF.P, and BD_BMA, can provide the edge existence probability or the inclusion probability. However, it needs to be clarified first that it is not appropriate to compare directly the probability values derived from BMRF and BD_BMA, since the models considered are different. In BMRF, the existence probability of the edge is the posterior probability of *γ_jk_* = 1; while in BD_BMA, it is the sum of all posterior probabilities of networks containing the edge, also called the inclusion probability. This quantity is also the expected value of its existence. Therefore, it would be reasonable if the edge probability in BD_BMA is slightly larger than that in BMRF. In other words, if the same threshold is used to determine whether an edge exists, the BMRF would be more conservative than BD_BMA. This can be observed in Figure 2 (c): for truly existent edges (Edge=1 on X-axis), these three methods all provide a large probability of existence, with that of BD_BMA being the largest; while if truly no edge exists (Edge=0 on X-axis), the three all provide very low probabilities. Note that this observation is consistent with the findings of TP and SEN in M1 and M2. Moreover, since BMRF.P utilizes more information than BMRF.O, in this case the information from SPACE, it is expected that BMRF.P will achieve a larger probability of existence than BMRF.O, as demonstrated in Figure 2 (c). Although it is not proper to compare the existence probability directly with the inclusion probability, we can examine their associations. The probabilities under BMRF.P versus that under BD_BMA are plotted in Figure 2 (d), where blue circles represent true edges and the red circles indicate non-existent edges. These two are fairly consistent, and BMRF.P performs better than BD_BMA.

**Figure 2.**
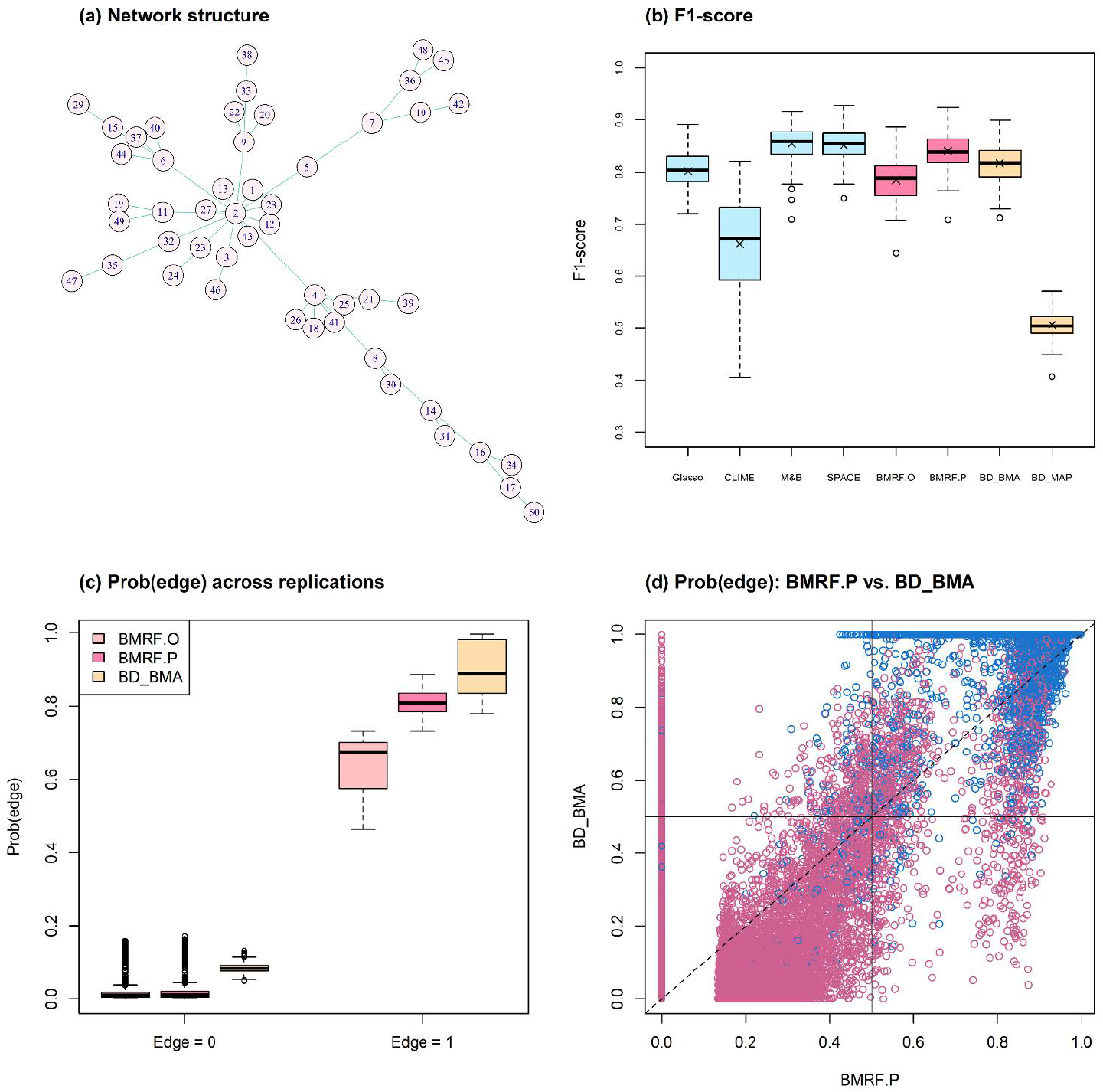
Simulation studies for setting M3. (a) The true network structure with 49 true edges; (b) Boxplots of the F1-score from 100 replications under each method; (c) Plots of average existence probability over 100 replications. The left panel (Edge = 0) is for the non-existent edges and the right panel (Edge=1) contains 49 average probabilities in each boxplot; (d) The edge existence probability from BMRF.P versus the inclusion probability from BD_BMA for each edge across replications. Blue circles indicate true edges and red indicates no edge. The vertical and horizontal solid lines denote the cut-off values for BMRF.P and BD_BMA, respectively.

**Table 2.**
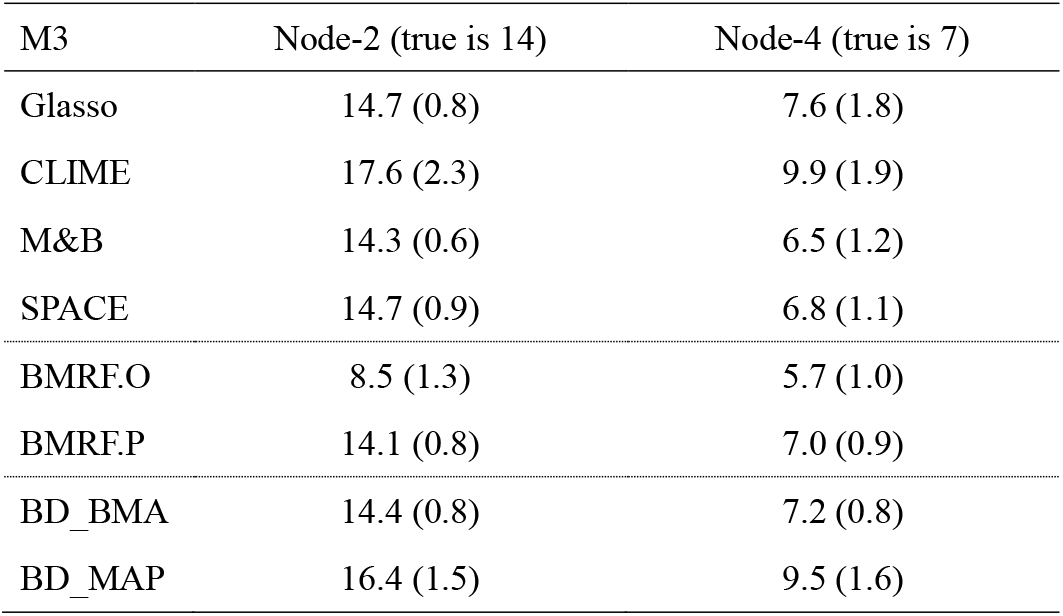
The listed values are the average number of estimated edges connecting to each of the two hub nodes (Node-2 and Node-4) across 100 replications under M3. The number in parenthesis is the standard error.

## 4. THE GLIOBLASTOMA STUDY

Glioblastoma (GBM) is a grade IV malignant brain tumor, usually in adults. After being diagnosed, patients have a median survival time of about 12 to 15 months and generally respond poorly to treatments (Stupp et al., 2005; TCGA, 2008). Although several molecular biomarkers have been identified, such as TP53 mutation and overexpression in EGFR (Bralten and French, 2011; Zhang et al., 2018), targeted therapy shows a limited effect (Shergalis et al., 2018; Banerjee et al., 2021). Recently, interest has focused on the molecular mechanism of the Janus kinase/signal transducer and activator of transcription (JAK-STAT) signaling pathway (Jain et al., 2012; Ou et al., 2021). Here, the BMRF model is applied to the JAK-STAT signaling pathway to examine the conditional dependence among gene nodes and detect influential molecular relationships to understand the underlying biological mechanism better.

The GBM dataset was downloaded from the University of California STANTA CRUZ (UCSC Xena) TCGA Hub. It contains the gene expression data generated from the Affymetrix HT Human Genome U133a microarray platform with mRNA values in the log 2 scale. The procedures in Chang et al. (2020) were then used to extract the information of genes in the JAK-STAT pathway. The final data consist of 27 gene expression values from 253 primary tumor tissues of male patients aged 40 and 75. The set of edges corresponding to the top 15% sample correlation and edges identified by SPACE was first determined, leading to a set containing 99 edges and a large sparsity of around 0.3. More information about the edges is in the Supporting Information Figures S1 and S2.

### 4.1 Edge identification

BMRF.P identified 69 edges, 15 of which were associated with a posterior existence probability greater than 0.9. Figure 3 (a) shows the resulting gene regulatory network, where the 15 edges are represented with thick lines and the others with thin lines. The corresponding magnitudes of the 15 existence probabilities are displayed in Figure 3 (b), where the width denotes the magnitude of the probability. The boxplots in Figure 3 (c) show the posterior samples of the strength of each edge, all displaying positive conditional correlations between paired nodes. This is consistent with the pattern of co-expression, and the first two pairs seem to be strongly correlated with each other. Other summary statistics regarding these 15 edges and all 69 edges are provided in the Supporting Information Table S4 and Figure S3, respectively. In Figure 4, the existence probabilities are ordered if prioritization is of interest.

**Figure 3.**
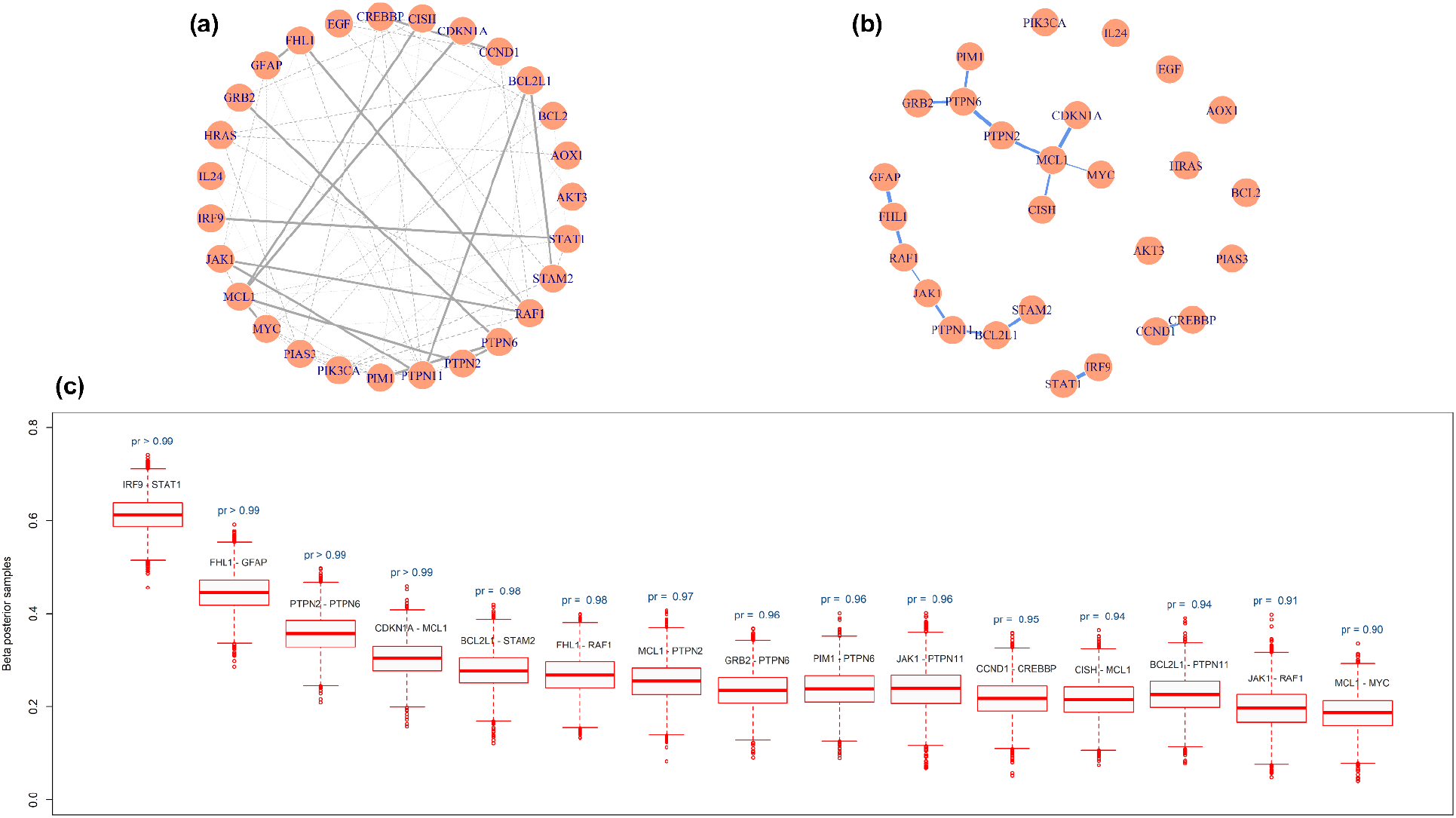
Gene regulatory network constructed by BMRF.P. (a) The estimated genetic network. The 15 thick lines are edges with an estimated existence probability greater than 0.9; (b) The network structure containing only the 15 edges, where the width of the edge corresponds to the magnitude of the existence probability; (c) Boxplots of the posterior samples of the strength coefficient corresponding to each one of the 15 edges. The text above the boxplot represents the estimated existence probability.

**Figure 4.**
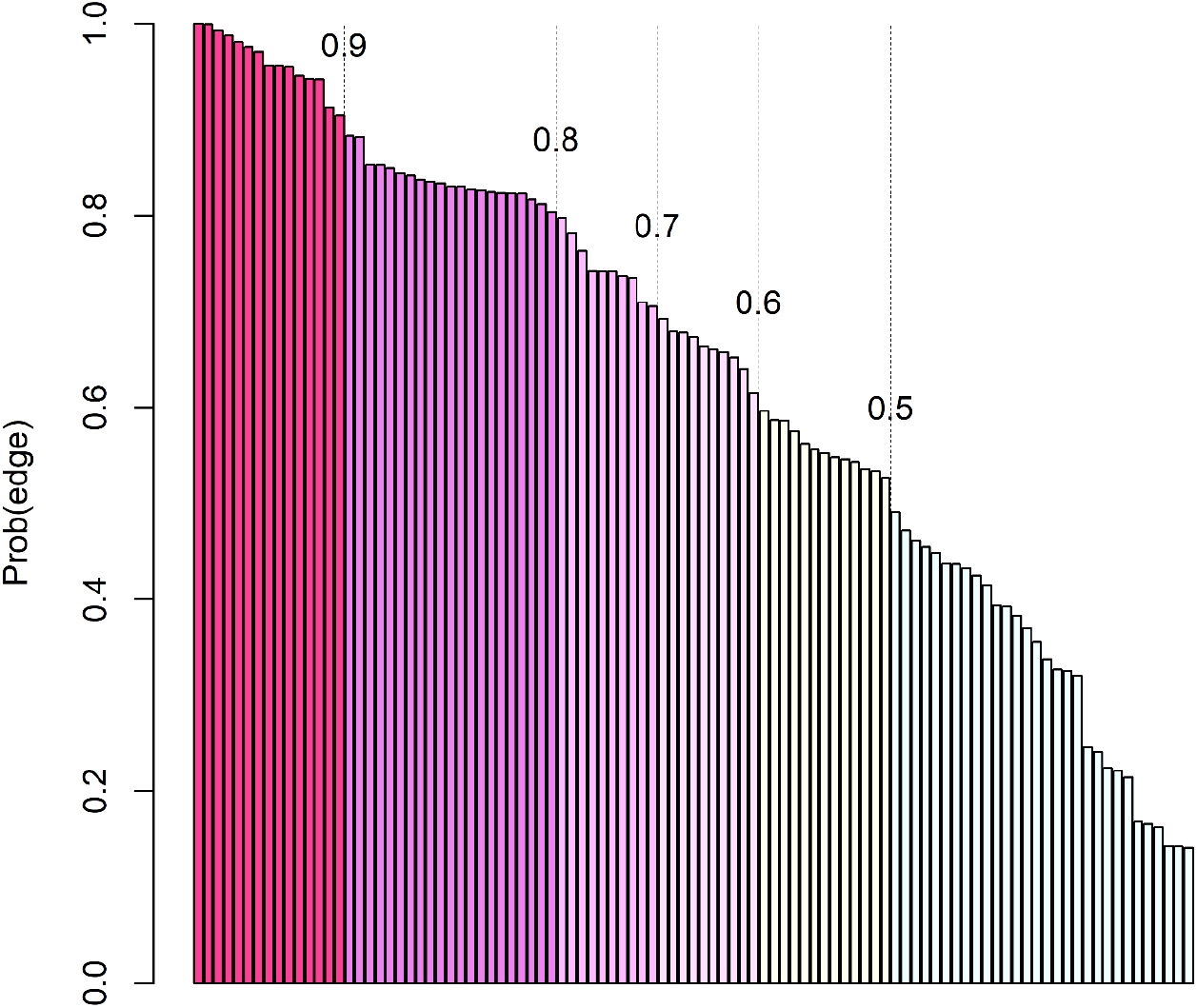
The ordered probabilities of the 99 edges are estimated by BMRF.P, and different colors correspond to different thresholds. The first 69 are the edges with a probability greater than 0.5.

In the Supporting Information Figures S4 and S5, the findings of BMRF.P are compared with those of alternative procedures. All edges identified by BMRF.P overlap with those identified by other procedures. Similar to the simulation studies, the edges identified by CLIME and BD_BMA overlap the least with the other procedures.

### 4.2 Biological interpretations

The proposed BMRF detected several influential biomarkers in the GBM study. First, the identified hub node *MCL1* in Figure 3 (b) has been reported as one of the cell apoptosis inhibitors associated with the progression of GMB and participates in the signaling of the maintenance of neural stem cells (Fassl et al., 2012; Murphy et al., 2014). In addition, the transcription factor c-Myc of the gene *MYC* connected to *MCL1* plays a central role in regulating the proliferation and survival of glioblastoma stem cells (Wang et al., 2008; Ha et al., 2015). Moreover, in the network constructed by BMRF.P, the genes (*PTPN2, PTPN6, PTPN11*) in the Protein-Tyrosine Phosphatase Non-Receptor (PTPN) family take critical positions. This is not surprising since the expression level of the immunotherapy target PTP2 has been shown to associate with the grade of glioma (Wang et al., 2018). Liu and colleagues (Liu et al., 2011) have suggested *PTPN11* as a functional target for treating glioblastomas in human and animal studies, and Cerami et al. (2010) have identified *PTPN11* as associated with an oncogenic process in GBM patients. Members of the PTPN family induce dephosphorylation of *JAK*, thereby regulating JAK-STAT signaling (Xu and Qu, 2008; Jain et al., 2012; Hammarén et al., 2019) In addition, the largest partial correlation was found between *IRF9* and *STAT1*. Au-Yeung et al. (2013) noted that this interaction is involved in type I interferon (IFN) signaling and anti-viral immune response. We have summarized other interactions in the Supporting Information Table S5.

## 5. CONCLUSION AND DISCUSSION

The BMRF approach introduced here provides the probabilistic inference of edges in a network graph, including the existence probability and the relative strength of edges. The proposed model starts with the conditional autoregressive model and SSL prior, and then establishes the posterior distribution of *β_jk_* to indicate the intensity of partial correlation and the marginal posterior probability of *γ_ij_*=1 to infer the edge existence probability. Simulation studies have demonstrated that the BMRF model can provide performance comparable with existing methods even when only the existence is of interest. For a network with a particular structure, such as the scale-free network, the performance of BMRF can be significantly improved if other prior information is incorporated. The glioblastoma study has demonstrated that the BMRF model can reproduce previous scientific findings and provide a list of prioritized pairwise conditional dependence for future biological or therapeutic experiments. The code in R can be downloaded from https://github.com/YJGene0806/BMRF_Code.

Although BMRF and BDgraph are both Bayesian approaches, the emphases are different. BMRF focuses on inference of the conditional relationship, while BDgraph is more interested in identifying non-zero elements in the precision matrix. Ideas similar to BMRF may be executed with other Bayesian models. For instance, Wang (2012), Peterson et al. (2013), and Gan et al. (2019) assigned for the precision matrix a prior distribution composed of a product of all probability distributions of each element. Though such an approach can estimate each edge directly, it is hard to evaluate the entire network’s uncertainty and track the precision matrix’s structure. Even the G-Wishart prior adopted in Wang and Li (2012) and BDgraph requires further post-processing techniques to achieve the probability of edge existence. This direction warrants further investigation.

The proposed BMRF can be extended to integrative network analysis. With a graphical model comprised of biomarkers from different platforms, it is possible to reveal the underlying complex biological structure among various forms of molecules (Peng et al., 2010; Yin and Li, 2011; Cai et al., 2013; Ha et al., 2021). In this case, adjustments in the CAR model would be needed to account for the genetic variables at different levels. However, this approach would be computationally intensive when facing the enormous number of all parameters combined.

Another generalization of the BMRF is to relax the distributional assumption in the CAR model. The GGM for the gene network assumes the MVN as the joint distribution, and the conditional and marginal distribution are also Gaussian. This assumption may not be valid generally (Ho et al., 2021). Classical research has addressed the non-Gaussian type of Markov random field (Besag, 1974), but these studies are not designed for sparse neighborhood selection. One solution would be to combine the non-paranormal distribution in Liu et al. (2009) or the exponential family graphical model (Yang et al., 2015) with BMRF for further investigation.

## ACKNOWLEDGMENT

This work was partially supported by MOST 109-2314-B-002-152 and MOST 110-2314-B-002-078-MY3.

## Web Appendix A: Supporting information for simulation studies

**Table S1.** Details of simulations. For example, in setting (M1.1), the average number of true edges across 100 replications is around 15.

| Setting |  | # of true edges | Sparsity |
| --- | --- | --- | --- |
| (M1.1) Random network | P = 25 | $\approx 15$ | 0.05 |
| (M1.2) Random network | P = 25 | $\approx 30$ | 0.10 |
| (M1.3) Random network | P = 50 | $\approx 62$ | 0.05 |
| (M1.4) Random network | P = 50 | $\approx 123$ | 0.10 |
| (M2.1) Scale-free network | P = 25 | 24 | 0.08 |
| (M2.2) Scale-free network | P = 50 | 49 | 0.04 |
| (M.3) Fixed network | P = 50 | 49 | 0.04 |

**Table S2.** Values of evaluation criteria under simulation settings M1.2, M1.4, and M2.1. Each value is the average of 100 replications with standard error (SE) in parentheses. The false discovery rate (FDR) is defined as the average value of FDP.

| M1.2 | F1 | MCC | FDR | TP | SEN | SPE |
| --- | --- | --- | --- | --- | --- | --- |
| Glasso | 0.80 (0.06) | 0.78 (0.06) | 0.18 (0.08) | 23.2 (3.1) | 0.78 (0.10) | 0.98 (0.010) |
| CLIME | 0.73 (0.11) | 0.74 (0.10) | 0.01 (0.03) | 17.4 (4.1) | 0.59 (0.10) | 0.99 (0.004) |
| M&B | 0.83 (0.06) | 0.82 (0.07) | 0.09 (0.05) | 22.8 (3.1) | 0.77 (0.10) | 0.99 (0.005) |
| SPACE | 0.83 (0.06) | 0.81 (0.07) | 0.10 (0.05) | 23.0 (3.2) | 0.77 (0.10) | 0.99 (0.006) |
| BMRF.O | 0.82 (0.07) | 0.81 (0.07) | 0.07 (0.05) | 22.1 (3.0) | 0.74 (0.10) | 0.99 (0.005) |
| BMRF.P | 0.83 (0.06) | 0.82 (0.06) | 0.11 (0.05) | 23.4 (3.1) | 0.78 (0.09) | 0.99 (0.006) |
| BD_BMA | 0.83 (0.06) | 0.82 (0.06) | 0.12 (0.06) | 23.7 (3.1) | 0.80 (0.09) | 0.99 (0.007) |
| BD_MAP | 0.65 (0.06) | 0.62 (0.07) | 0.45 (0.07) | 23.8 (3.1) | 0.80 (0.09) | 0.93 (0.020) |
| M1.4 | F1 | MCC | FDR | TP | SEN | SPE |
| Glasso | 0.43 (0.07) | 0.49 (0.06) | 0.06 (0.05) | 34.5 (6.1) | 0.28 (0.07) | 0.99 (0.002) |
| CLIME | 0.47 (0.09) | 0.50 (0.06) | 0.12 (0.10) | 41.0 (12.2) | 0.33 (0.10) | 0.99 (0.009) |
| M&B | 0.43 (0.07) | 0.49 (0.05) | 0.06 (0.04) | 34.2 (5.4) | 0.28 (0.06) | 0.99 (0.002) |
| SPACE | 0.51 (0.07) | 0.53 (0.05) | 0.12 (0.05) | 44.1 (5.1) | 0.36 (0.06) | 0.99 (0.003) |
| BMRF.O | 0.53 (0.05) | 0.54 (0.05) | 0.17 (0.05) | 48.0 (4.0) | 0.39 (0.05) | 0.99 (0.003) |
| BMRF.P | 0.54 (0.05) | 0.55 (0.05) | 0.19 (0.05) | 50.3 (4.6) | 0.41 (0.06) | 0.99 (0.003) |
| BD_BMA | 0.56 (0.05) | 0.55 (0.05) | 0.23 (0.05) | 54.0 (4.0) | 0.44 (0.05) | 0.99 (0.004) |
| BD_MAP | 0.45 (0.03) | 0.38 (0.04) | 0.57 (0.03) | 58.2 (4.3) | 0.47 (0.05) | 0.93 (0.006) |
| M2.1 | F1 | MCC | FDR | TP | SEN | SPE |
| Glasso | 0.81 (0.06) | 0.80 (0.06) | 0.31 (0.09) | 23.5 (0.7) | 0.98 (0.03) | 0.96 (0.017) |
| CLIME | 0.75 (0.09) | 0.75 (0.09) | 0.39 (0.12) | 23.8 (0.5) | 0.99 (0.02) | 0.94 (0.033) |
| M&B | 0.93 (0.04) | 0.92 (0.05) | 0.12 (0.07) | 23.6 (0.7) | 0.98 (0.03) | 0.99 (0.008) |
| SPACE | 0.90 (0.04) | 0.90 (0.04) | 0.16 (0.06) | 23.6 (0.7) | 0.98 (0.03) | 0.98 (0.008) |
| BMRF.O | 0.90 (0.08) | 0.89 (0.08) | 0.11 (0.07) | 21.8 (2.2) | 0.91 (0.09) | 0.99 (0.007) |
| BMRF.P | 0.90 (0.04) | 0.90 (0.04) | 0.16 (0.07) | 23.6 (0.6) | 0.98 (0.03) | 0.98 (0.008) |
| BD_BMA | 0.92 (0.04) | 0.92 (0.04) | 0.13 (0.06) | 23.7 (0.5) | 0.99 (0.02) | 0.99 (0.007) |
| BD_MAP | 0.72 (0.06) | 0.72 (0.06) | 0.43 (0.07) | 23.5 (0.7) | 0.98 (0.03) | 0.93 (0.019) |

**Table S3.**
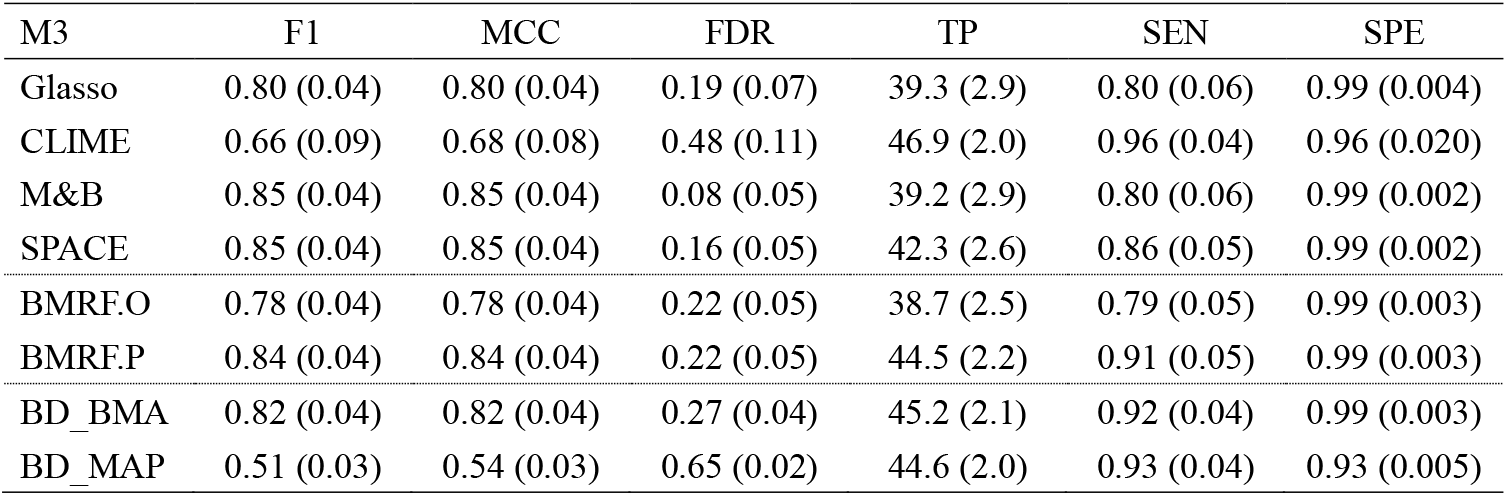
Values of evaluation criteria under simulation setting M3. Each value is the average of 100 replications with standard error (SE) in parentheses. The false discovery rate (FDR) is defined as the average value of FDP.

| M3 | F1 | MCC | FDR | TP | SEN | SPE |
| --- | --- | --- | --- | --- | --- | --- |
| Glasso | 0.80 (0.04) | 0.80 (0.04) | 0.19 (0.07) | 39.3 (2.9) | 0.80 (0.06) | 0.99 (0.004) |
| CLIME | 0.66 (0.09) | 0.68 (0.08) | 0.48 (0.11) | 46.9 (2.0) | 0.96 (0.04) | 0.96 (0.020) |
| M&B | 0.85 (0.04) | 0.85 (0.04) | 0.08 (0.05) | 39.2 (2.9) | 0.80 (0.06) | 0.99 (0.002) |
| SPACE | 0.85 (0.04) | 0.85 (0.04) | 0.16 (0.05) | 42.3 (2.6) | 0.86 (0.05) | 0.99 (0.002) |
| BMRF.O | 0.78 (0.04) | 0.78 (0.04) | 0.22 (0.05) | 38.7 (2.5) | 0.79 (0.05) | 0.99 (0.003) |
| BMRF.P | 0.84 (0.04) | 0.84 (0.04) | 0.22 (0.05) | 44.5 (2.2) | 0.91 (0.05) | 0.99 (0.003) |
| BD_BMA | 0.82 (0.04) | 0.82 (0.04) | 0.27 (0.04) | 45.2 (2.1) | 0.92 (0.04) | 0.99 (0.003) |
| BD_MAP | 0.51 (0.03) | 0.54 (0.03) | 0.65 (0.02) | 44.6 (2.0) | 0.93 (0.04) | 0.93 (0.005) |

## Web Appendix B: Supporting information for the TCGA glioblastoma study

**Figure S1.**
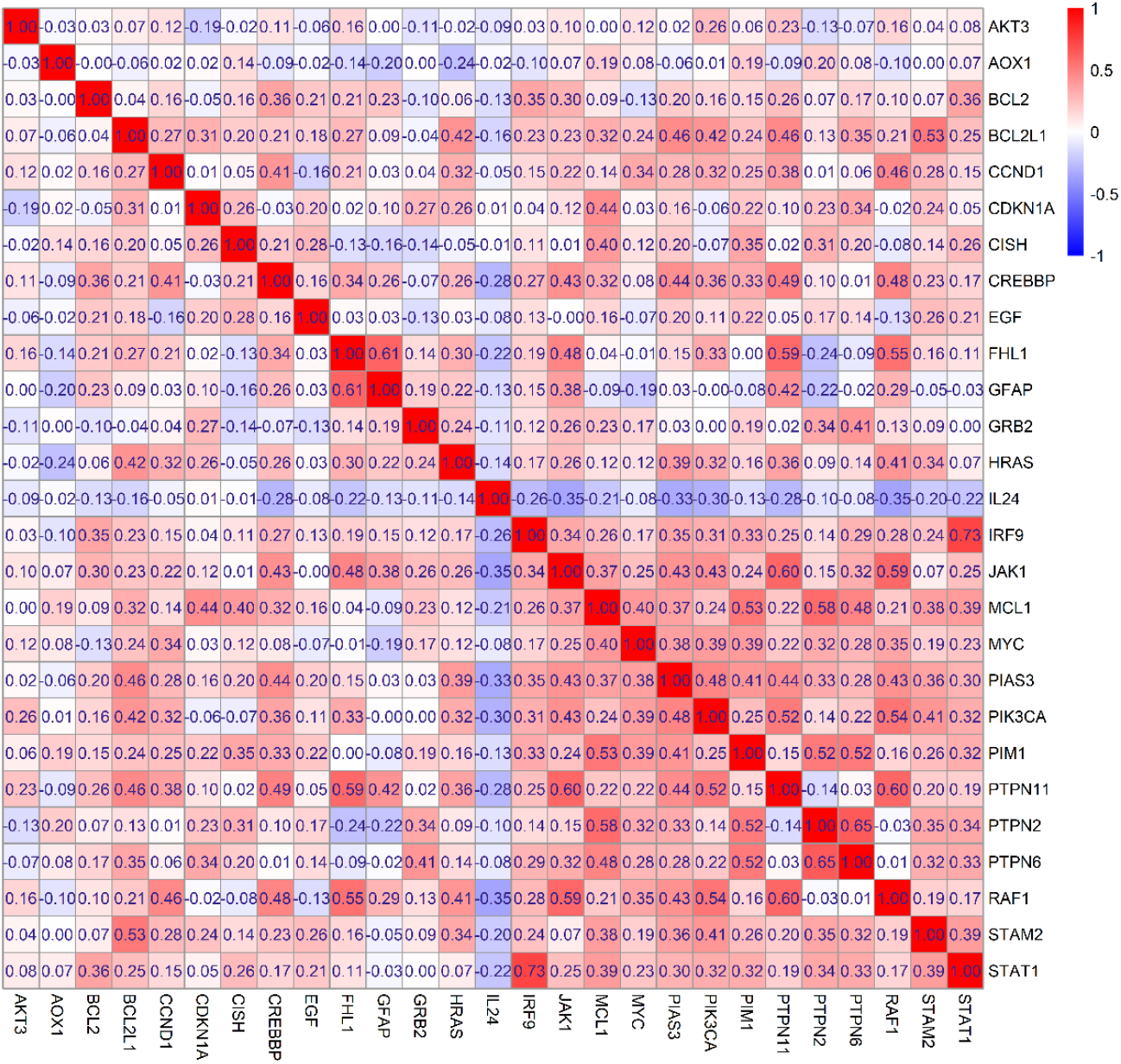
Sample correlation matrix.

**Figure S2.**
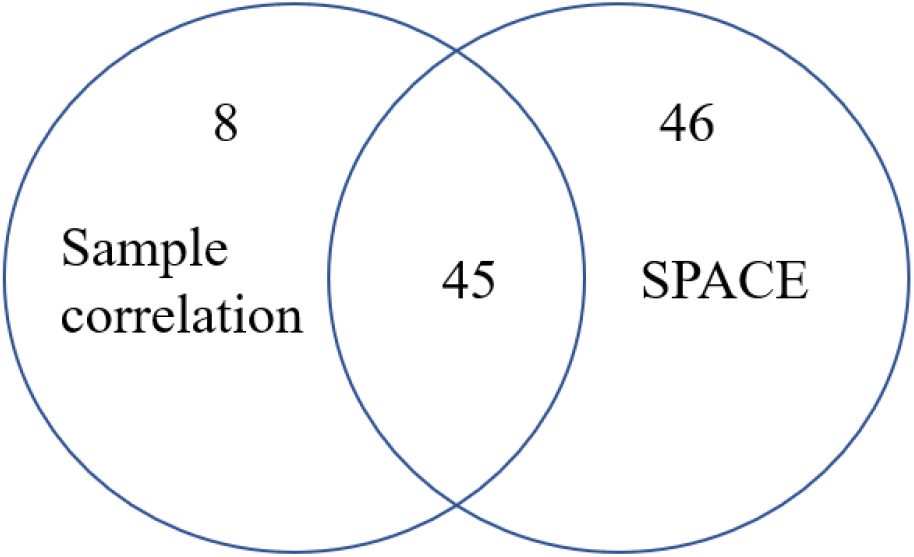
Venn diagram of the number of edges passing sample correlation screening and identified by SPACE in the GBM study. SPACE identified 91 edges and sample correlation screened the top 53 edges 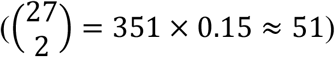. Forty-five edges are in common.

**Figure S3.**
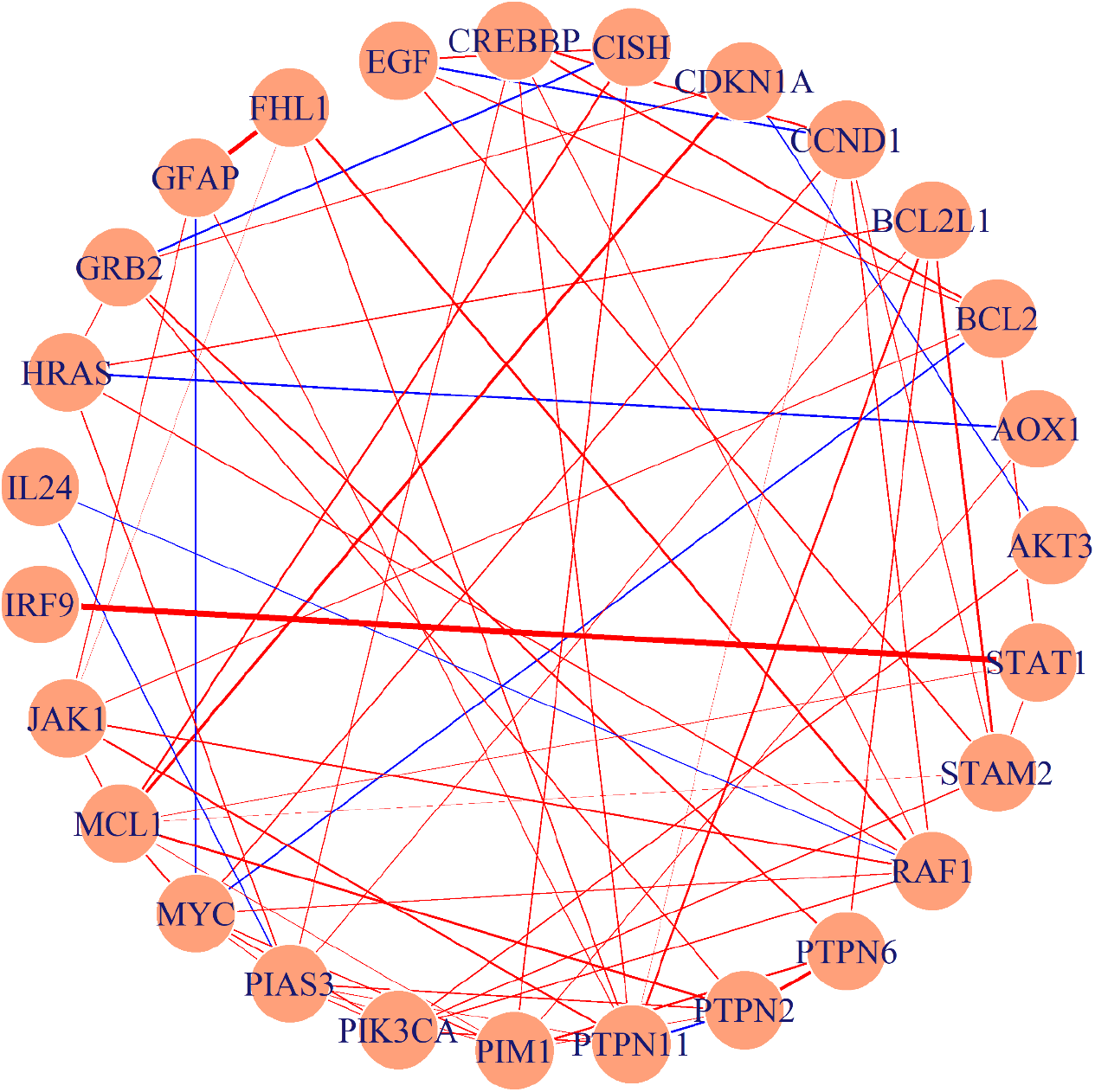
The gene regulatory network constructed by BMRF.P. The width of each edge represents the magnitude of the posterior mean of 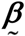. Color in red (blue) represents the positive (negative) partial correlation.

**Table S4.**
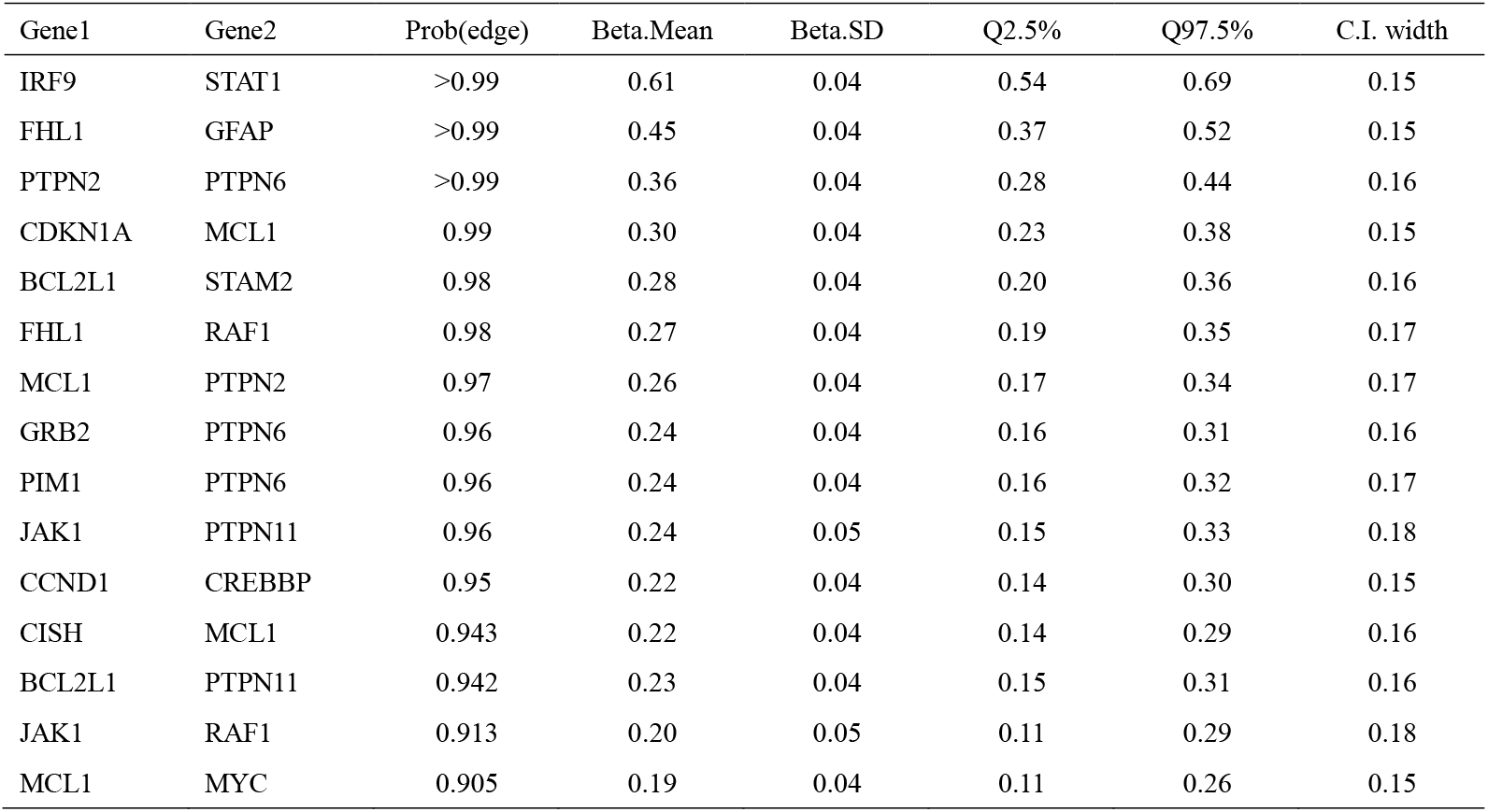
Details about the 15 edges (existence probability >0.9). (1) Gene1, Gene2: the two genes connected by the selected edge (2) Prob(edge): the estimated probability of existence (3) Beta.mean: the posterior mean (4) Beta.SD: the standard deviation of the beta posterior samples (5) Q2.5%, Q97.5%, and C.I. width: the quantiles and width of the 95% credible interval of beta

**Figure S4.**
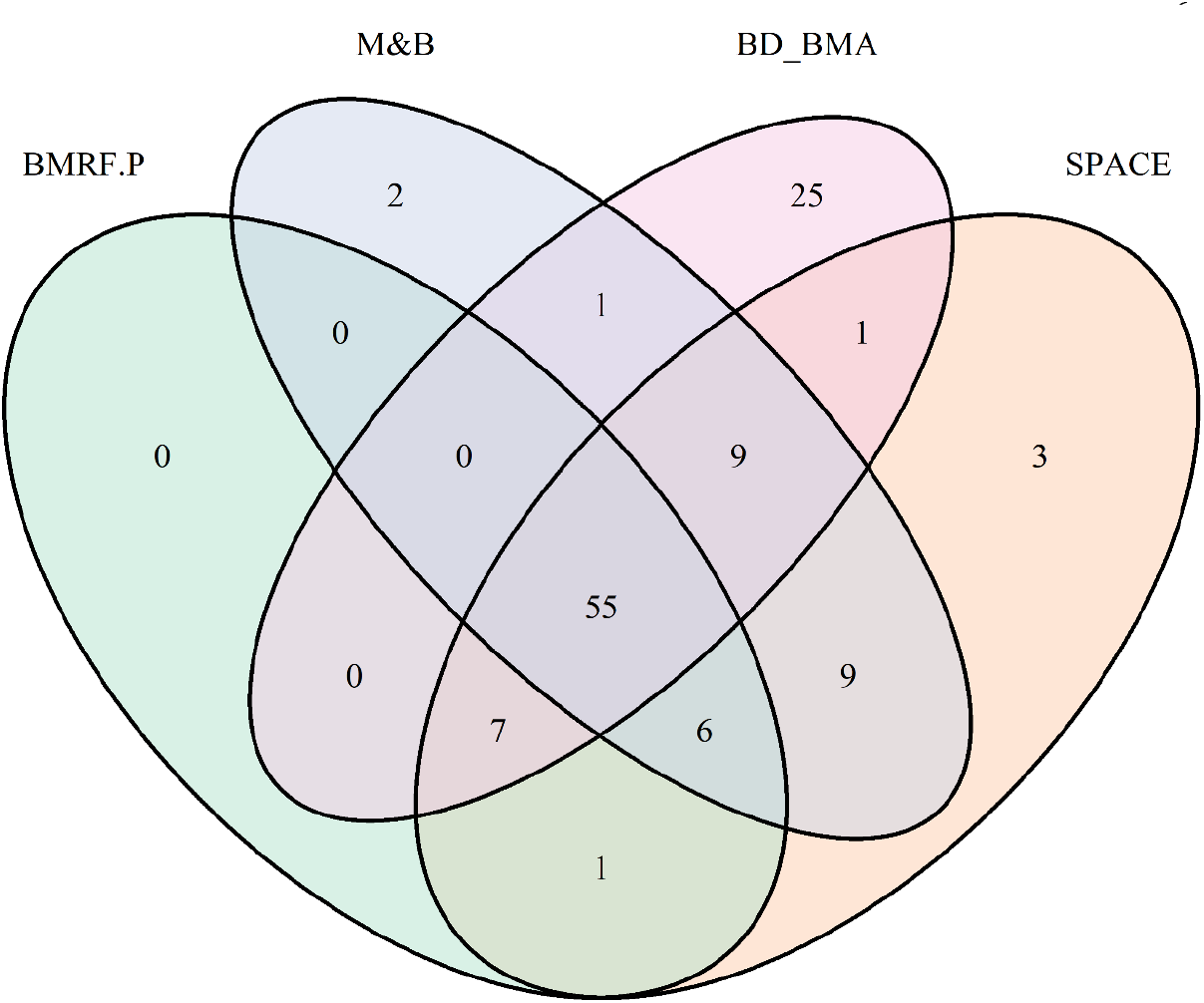
Venn diagram of the number of edges identified by BMRF.P, M&B, BD_BMA, and SPACE.

**Figure S5.**
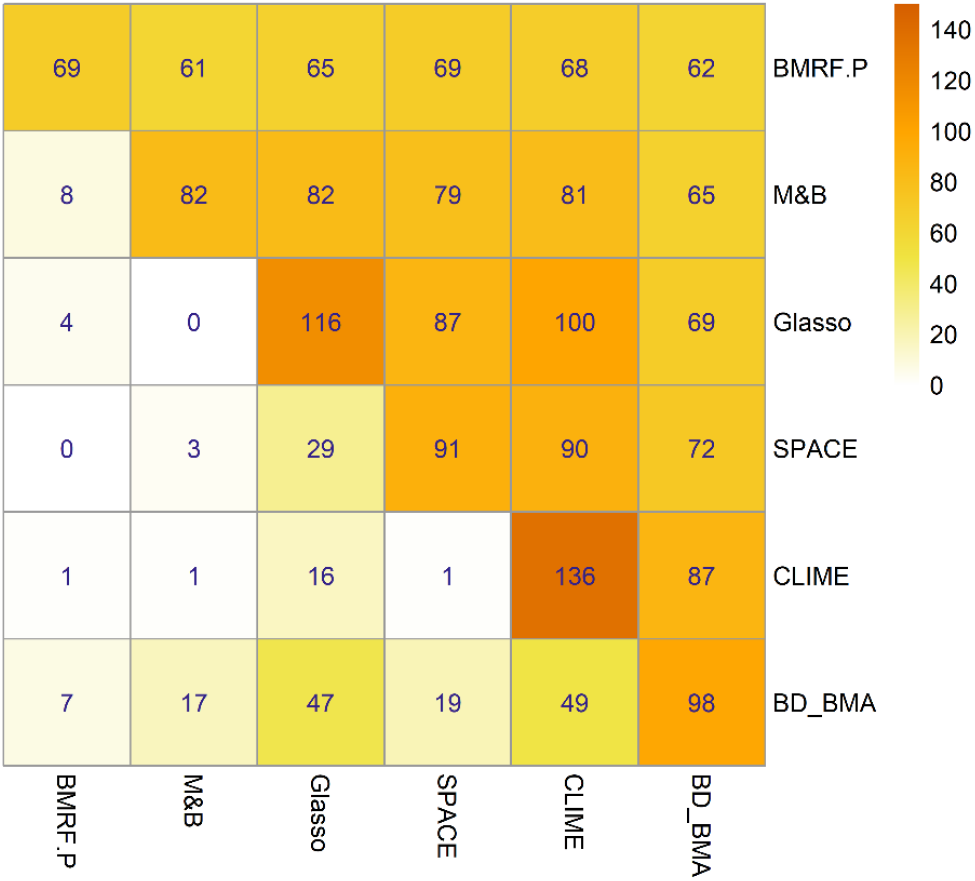
Numbers of identified edges. (1) Numbers in the diagonal represent the number of edges identified by each method. (2) Numbers in the upper triangle represent the number of edges identified by both the column and row methods. (3) Numbers in the lower triangle represent the number of edges identified by the column method but not the row method. For example, BMRF.P has identified 69 edges. Among them, 61 edges have been identified by M&B as well, while 8 among the 69 were identified by BMRF.P and not by M&B.

**Table S5.** List of selective interactions that have been reported in public databases. The symbol “✓” indicates the edge was successfully identified by the corresponding method, and the symbol “✗” indicates failure to identify.

| Gene1 | Gene2 | BMRF.P<br>(prob. of<br>existence) | BMRF.P<br>(95% C.I. of<br>beta) | M&B | Glasso | SPACE | CLIME | BD_BMA | KEGG | String | BioGrid |
| --- | --- | --- | --- | --- | --- | --- | --- | --- | --- | --- | --- |
| IRF9 | STAT1 | >0.99 (✓) | (0.54, 0.69) | ✓ | ✓ | ✓ | ✓ | >0.99 (✓) | ✓ | ✓ | ✓ |
| JAK1 | PTPN11 | 0.96 (✓) | (0.15, 0.33) | ✓ | ✓ | ✓ | ✓ | 0.99 (✓) | ✓ | ✓ | ✓ |
| RAF1 | HRAS | 0.83 (✓) | (0.07, 0.23) | ✓ | ✓ | ✓ | ✓ | >0.99 (✓) | ✓ | ✓ | ✓ |
| MYC | BLC2 | 0.80 (✓) | (-0.28, -0.13) | ✗ | ✗ | ✓ | ✓ | 0.92 (✓) | ✗ | ✗ | ✓ |

